# Oscillatory brain activity reflects semantic and phonological activation during sentence planning

**DOI:** 10.1101/2025.11.26.690808

**Authors:** Jed A. Meltzer, Aneta Kielar, Frank Oppermann

## Abstract

Verbal short-term memory includes resources for maintaining semantic and phonological information. These resources are complementary and often activated simultaneously, making their anatomical bases difficult to determine. One way to distinguish them may be to study the resolution of interference from distractor words that are semantically or phonologically related to a planned sentence. We recorded magnetoencephalography (MEG) data while participants rehearsed short formulaic 5-word sentences like “The mouse ate the cheese.” During a memory delay period, participants exhibited bilateral temporofrontal event-related desynchronization (ERD, power decrease) in the alpha and beta bands (8-30 Hz). During the memory delay, participants also heard an auditory distractor word that could be unrelated to the sentence, semantically related to one of the words in the rehearsed sentence (e.g. “rat” or “butter”), or phonologically related to one of the words in the sentence (e.g. “mountain” or “cheat”). Relative to unrelated words, related words induced a greater degree of ERD immediately following their presentation. Effects of semantic distractors were exclusively in the temporal lobe, largely in the left middle temporal gyrus but also in bilateral medial temporal regions. Effects of phonological distractors were far more widespread in temporal, frontal, and parietal regions, and were largely left-lateralized, although they also overlapped with the temporal regions showing semantic effects. As no behavioural effects were observed in cued sentence repetition, it seems that auditory distractors produce short-lasting interference with a verbal memory trace that is ultimately resolved, but useful for mapping regions involved in maintaining distinct aspects of the sentence content.

## Introduction

The neural bases of language and working memory have much in common. Individual differences in verbal working memory capacity predict comprehension of syntactically complex sentences in both auditory and written modalities (Caplan and Waters, 1999), as well as general reading comprehension (Daneman and Carpenter, 1980). Increasing the difficulty of language comprehension tasks induces similar patterns of brain activation as increasing working memory load (Deldar et al., 2021; Emch et al., 2019). Researchers have distinguished multiple mechanisms including phonological short-term memory (pSTM), underlying “rote rehearsal,” and conceptual short-term memory (cSTM), a semantic resource that can support the regeneration of a sentence’s form from basic elements of its meaning (Potter and Lombardi, 1990; Little et al., 2006). Neuroimaging and brain stimulation studies have suggested a neuroanatomical dissociation between the two, with pSTM relying on fronto-parietal circuits, including more dorsal portions of the inferior frontal gyrus (IFG) (Buchsbaum and D’esposito, 2008; Klaus and Hartwigsen, 2019) while cSTM may depend on more ventrally located networks in the temporal lobe and lower portions of the IFG (Bonhage et al., 2014; Perrone-Bertolotti et al., 2017).

The role of working memory in spontaneous language production is less well characterized. When producing continuous speech, speakers must retrieve and maintain semantic and phonological information about the words that they intend to produce next. Studies of single-word production suggest that semantic retrieval precedes phonological retrieval in time, and that these processes respect a ventral/dorsal anatomical distinction with semantic retrieval occurring largely in the left temporal lobe, and phonological retrieval depending on a more extensive fronto-parietal network (Indefrey, 2011). However, it is uncertain whether similar distinctions exist in the mechanisms that account for advance planning of multi-word utterances.

Early evidence for a distinction between semantic and phonological aspects of speech planning came from analyses of speech error corpora, especially on transposition errors in which two words or sounds change places. According to a spreading activation assumption of word retrieval processes, these exchange errors occur when an upcoming unit of the utterance becomes more strongly activated at the same time as the current unit is selected (Dell, 1986). Interestingly, these errors have a different scope for lexical units and speech sounds (Garrett, 1975, 1980). Semantically related words are often exchanged between noun phrases with other words coming in between them (e.g. “put the lock in the key”), maintaining their proper syntactic function, whereas sounds are often exchanged between neighboring words irrespective of syntactic categories (e.g. “heft lemisphere” instead of “left hemisphere”). This suggests that the temporal scope for semantic and phonological representations may differ, with planning of semantic content extending further forward in time than planning of phonological content does.

Investigations of word retrieval have made extensive use of “Picture Word Interference” (PWI) tasks. In PWI, a participant is presented with a picture to be named, along with a distractor word, either written or spoken. On some trials, the distractor word is unrelated to the picture, while on others, it may be either semantically related (e.g. DOG → CAT) or phonologically related (e.g. BAT → BAG). Depending on the stimulus onset asynchrony (SOA) between the picture and the distractor, semantically or phonologically related words may exert facilitative effects, reducing reaction time and improving accuracy compared to unrelated words, or they may instead exert interference effects, resulting in increased reaction time and reduced accuracy (Schriefers et al., 1990; Burki et al., 2020). Most commonly, semantic distractors produce interference, especially when they come from the same semantic category (e.g. DOG → CAT but not DOG → BONE) (Abel et al., 2009). Semantic interference is thought to arise because the distractor word activates overlapping but competing semantic representations with the intended target, and the participant must resolve the competition (Roelofs, 1992). Phonological distractors, in contrast, frequently induce facilitation. For example, if one is to produce the word “bat,” then pre-activation of the first two phonemes through hearing the word “bag” may produce a net gain in neural efficiency (Starreveld and La Heij, 1995). Semantic interference occurs optimally when the distractor word precedes the target picture (e.g. SOA = −200 ms), whereas phonological facilitation occurs when the distractor word is presented simultaneously with the picture or slightly later (e.g. SOA = +100 ms), reflecting the fact that lexical retrieval comprises sequential (albeit possibly overlapping) stages of semantic retrieval followed by phonological form retrieval (Levelt, 2001). Although resolution of interference may not reflect exactly the same process as ordinary retrieval of semantic and phonological information related to word production, the results of PWI experiments map neatly onto the theoretical stages of word retrieval (Levelt, 2001), suggesting that probing the language production system with different forms of interference can be an effective means of dissociating mechanisms involved in retrieval and maintenance of word forms. In the present study, we apply this principle to electrophysiological responses to distractor words while participants maintain sentences in short-term memory, rather than to their behaviour at the time of speaking.

To investigate differences in temporal scope, researchers have developed a number of extensions of the PWI technique for multi-word utterances, typically presenting an auditory distractor word as participants attempt to initiate production of phrases or sentences, such as “The mouse ate the cheese.” The distractor word may then be semantically or phonologically related to an earlier or later word in the target sentence (see Table 1). Studies have yielded fairly divergent results. For example, Meyer (1996) found that the degree of inhibition induced by semantic distractors remained relatively constant across positions of the related word (Meyer, 1996), while Zhao and Yang (2006) found electrophysiological evidence that semantic interference was higher for earlier words. For phonological distractors, one study found facilitation for early words changing to no effect or even inhibition when the distractor is related to words coming later in the utterance (Oppermann et al., 2010), while another found facilitation at early and late positions (Schnur et al., 2006). Taken together, these divergent findings suggest that the scope of phonological and semantic planning may differ, but it is difficult to predict the direction of the difference. Furthermore, phonological advance planning may be more highly dependent on working memory resources, given a study that showed that its temporal scope, assessed by the techniques described above, is reduced under dual-task conditions with a verbal working memory load, whereas semantic planning remains intact (Klaus et al., 2017).

**TABLE 1:** Example distractor words for the sentence “The mouse ate the cheese.” SEM 1,2 = semantically related to first and second noun; PHO 1,2 = phonologically related to first and second noun. Unrelated words in this example served as related words for other experimental sentences, such that the exact same words appeared as both related and unrelated across trials.

| Example distractors for “The mouse ate the cheese.” |  |  |
| --- | --- | --- |
|  | **Related** | **Unrelated** |
| **SEM 1** | rat | firm |
| **SEM 2** | butter | rock |
| **PHO 1** | mountain | pine |
| **PHO 2** | chieftain | bowler |

In the present study, we used magnetoencephalography (MEG) to characterize the neural mechanisms that are involved in semantic and phonological, using a specialized sentence repetition task. On each trial, participants first read a 5-word sentence presented word for word on a screen, and then prepared to repeat it from memory. Sentences consisted of a two nouns and a verb, with appropriate determiners (e.g. “The mouse ate the cheese.”) Participants were instructed to withhold their verbal response until a visual cue appeared. During the delay, participants heard an auditory distractor word during the maintenance interval, which was either unrelated, or semantically or phonologically related to either the first or second noun in the sentence. Participants were instructed to simply ignore the distractor word and wait for the cue to recite the sentence.

Note that unlike studies of PWI, variance in the behaviour of speech production is not of primary interest here. Rather, the experiment investigated changes in the brain’s electrophysiological response to the distractor word. Auditory words are known to induce robust electrophysiological responses in widespread areas of cortex (Hari, 1991; Pulvermüller et al., 2006), including left-lateralized event-related desynchronization in the alpha and beta bands as seen in the present study (Youssofzadeh et al., 2020; Kim and Chung, 2008). We reasoned that relationships between the distractor words and the content of the sentences were likely to modulate neural responses in areas where neural activity helps to maintain such activation. For example, if a brain region responded differentially to semantically related vs. unrelated words, but not to phonologically related vs. unrelated words, this could indicate involvement in the maintenance of semantic but not phonological information in planned utterances. By measuring MEG responses to auditory distractor words that are semantically or phonologically related to words occurring early or late in the sentence, and comparing them to unrelated distractors, we sought to answer three outstanding questions about the neural mechanisms responsible for advance planning of utterances:

### 1 Localization of semantic vs. phonological representations

Based on prior studies (Majerus, 2013; Hickok and Poeppel, 2004), we believe that semantic information is maintained by ongoing activity in the ventral language network, mainly in the temporal lobe. In contrast, phonological information is represented in activity in more dorsal frontal-parietal regions. Thus, we hypothesized that semantically and phonologically related words would elicit modulated responses in these two networks, respectively. We further expected that activation within these networks would be left-lateralized, given the dominant role of the left hemisphere in language processing. Nonetheless, differences in lateralization are also of interest. The dual-stream model of language processing posits that semantic processing occurs within ventral structures bilaterally, but phonological speech planning is thought to be heavily left-lateralized (Hickok and Poeppel, 2004; 2007). On the other hand, a previous MEG study from our group showed increased engagement of right fronto-parietal regions when phonological maintenance demands were increased (Meltzer et al., 2017). Thus, we may expect differential responses to both semantic and phonological relatedness to occur bilaterally to some extent.

### 2 Scope of semantic vs. phonological planning

Based on prior studies with similar stimuli finding a more limited scope of phonological planning (Oppermann et al., 2010), we hypothesized that neural responses would be modulated by words semantically related to either the first or second noun in the sentence indiscriminately, but phonological relatedness effects would be more restricted to the first noun. However, it is possible that the present design may not be sensitive enough to detect such effects, because participants are rehearsing the sentence over a fairly long delay, and we cannot control what position they are at within the sentence while rehearsing it when the distractor word is heard.

### 3 Facilitation vs. interference

Neuroimaging studies have often shown that conditions that lead to behavioural facilitation also induce diminished neural responses, reflecting the fact that fewer resources are required to process a stimulus under conditions of facilitative pre-activation of relevant representations. On the other hand, conditions inducing competition, or inhibition, generally induce larger neural responses, reflecting the extra effort needed to resolve the competition. In imaging studies on picture-word interference, considerable variability has been reported on the directionality of effects for related vs. unrelated distractors, both with and without the presence of behavioural effects (for review see Arrigoni et al., 2025; de Zubicaray and Piai, 2019). We hypothesized that semantic distractors, inducing competition, would induce larger neural responses, whereas phonological distractors, which may facilitate utterance production, would induce smaller responses. On the other hand, it is possible that all effects could be interfering, since all words in the sentence are already “retrieved” at the time of distractor presentation, and so the net effect of any relatedness may be to disturb the rehearsal by activating representations of competitor words.

To assess neural activity related to maintenance of sentence content and processing of the distractor words, we applied two analyses of MEG data in source space. First, we measured changes in oscillatory power, focusing on event-related desynchronization (ERD) in the 8-30 Hz range, encompassing the alpha and beta bands of traditional EEG analysis. In several prior studies, we have observed that processing of linguistic stimuli and memory maintenance of linguistic information reliably induce this signal, with little distinction between alpha and beta (Meltzer et al., 2011; 2017; Kielar et al., 2015). We hypothesized that both maintenance of sentences and processing the distractor would induce 8-30 Hz ERD relative to baseline, and furthermore, that the degree of 8-30 Hz ERD would be modulated by relatedness. Second, we analyzed event-related fields, or time-domain averages of signals time-locked to auditory word onset. Prior studies with EEG have shown that both semantic and phonological auditory primes can modulate these signals, especially in the time range of 500-1000 ms post-stimulus (de Zubicaray and Piai, 2019). We anticipated that analyses of both signals would converge on similar conclusions.

## METHODS

### Participants

Twenty right-handed, healthy young adults participated in the experiment (8 men; mean age 24.1 years, s.d. 2.231). Participants were recruited through advertisements from the Greater Toronto Area and the University of Toronto community. All were monolingual native speakers of English, with normal hearing and normal or corrected-to-normal vision. None had a history of neurological or psychiatric illness, had experienced neurological injury, or had used psychotropic medication. Participants gave informed consent and were financially compensated for their time.

### Procedure

Participants were seated in a padded chair inside a magnetically shielded room containing the MEG instrument. Visual stimuli were presented on a screen approximately one meter in front of the participant’s face. Auditory stimuli were delivered through pneumatic tubes ending in foam-insert earphones, with the volume individually adjusted so that the participant could hear the words comfortably. Participants’ auditory responses were recorded using an MEG-compatible fibreoptic microphone (Optoacoustics, Moshav Mazor, Israel).

The sentence repetition paradigm (Figure 1) was programmed using Presentation software (Neurobehavioral Systems, Berkeley, CA, USA). Each trial began with a fixation cross presented for 300 ms, followed by a blank screen for 200 ms. Next, the visual sentence was presented in RSVP format, with each word presented for 50 ms at a rate of one word per 200 ms. After the sentence was complete, a pattern mask (%%%%%%%) was presented for 500 ms. All sentences were simple declarative past tense statements with two nouns (subject and object) and a verb. In subsequent sentences, we will refer to the two nouns as noun 1 and 2, simply reflecting their chronological order in the sentence. Most sentences consisted of five words (e.g. “The mouse ate the cheese”) although some had six words with a preposition included in the verb phrase (e.g. “The pilot asked for a bonus.”) To prevent participants from invariably anticipating the auditory prime word, one ninth of trials (11.1%) were “catch trials” without a prime word. In catch trials, the cue to begin reciting the sentence (“!!!”) appeared 1,100 ms after the offset of the mask of the last word in the sentence. On all other trials, the recitation cue appeared 2,600 ms after the mask offset, and the auditory prime word was presented at a jittered interval 900-1,100 ms after the mask offset. The auditory prime word could be semantically or phonologically related to noun 1 or noun 2, but such relationships were irrelevant to the participant; they were instructed to ignore the distractor word and to attempt to recite the sentence that they had read verbatim. The recitation cue remained on the screen for 1000 ms and was followed by 2000 ms of a blank screen before the onset of a fixation cross alerting the participant to the beginning of the next trial. The stimulus presentation software automatically recorded the participant’s speech for a period of 5000 ms after each recitation cue.

**Figure 1.**
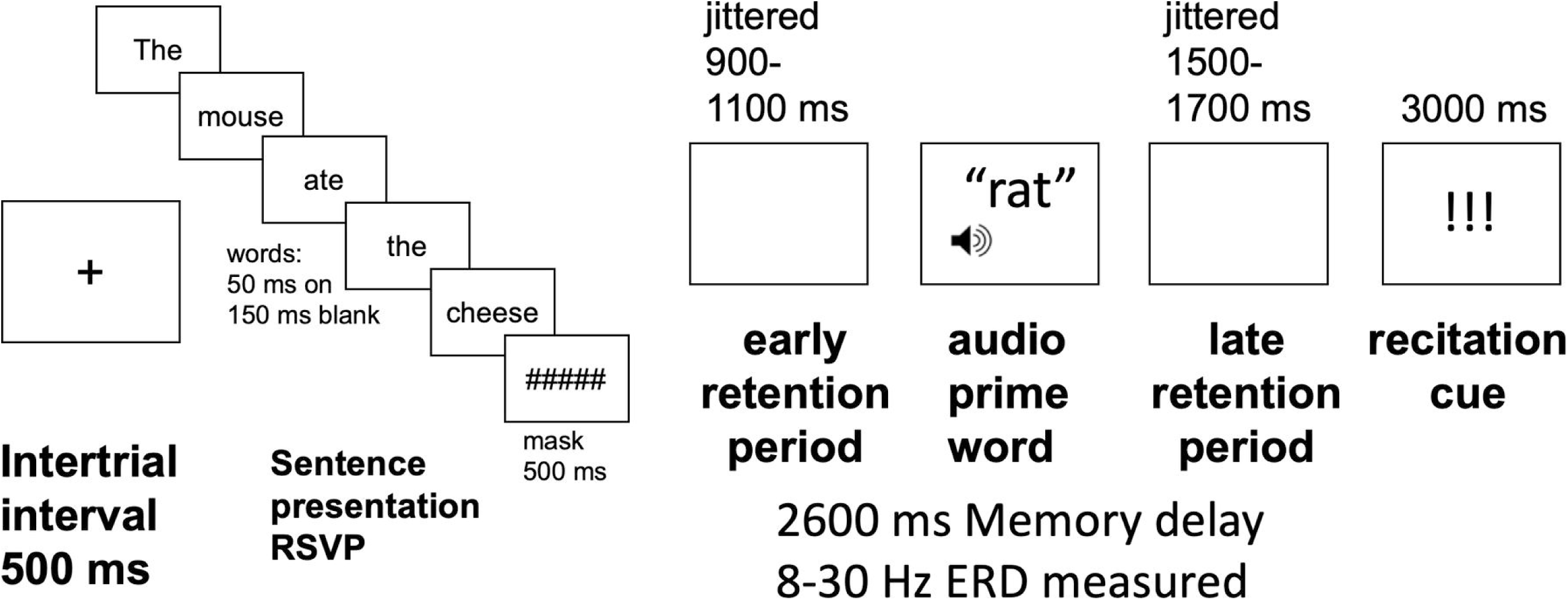
Task and trial design.

### Materials design

48 sentences were created for the experiment, and each was presented 9 times over the course of the experiment (1 catch trial plus 8 experimental conditions). A separate set of 8 sentences was used for practice trials to instruct the participants on the task.

Auditory prime words were digitally recorded by a female English speaker, using a consistent upward pitch intonation. Each prime word was either related to one word in the sentence, or unrelated to all of the words. If the word was related, it could be either related to noun 1 or to noun 2, and the relationship could be either semantic or phonological. Thus, 4 conditions of relatedness were included:

Sem1rel - semantically related to the first noun.
Sem2rel - semantically related to the second noun.
Pho1rel - phonologically related to the first noun.
Pho2rel - phonologically related to the second noun.

Additional words were chosen for the unrelated control conditions.

Sem1unr - semantically unrelated to the first noun.
Sem2unr - semantically unrelated to the second noun.
Pho1unr - phonologically unrelated to the first noun.
Pho2unr - phonologically unrelated to the second noun.

Example distractor words in these different conditions are shown in Table 1. With 48 experimental sentences, each with a two nouns, this resulted in 96 words to be paired with a semantic prime, a phonological prime, and two unrelated distractor words. 192 distractor words were chosen for the experiment, with each one being used twice: once as a semantically or phonologically related prime for a word in the experimental sentences, and once as an unrelated control word paired with a different sentence. Thus, all comparisons of related vs. unrelated words *involved exactly the same distractor words being heard*, with the context of the sentence determining whether they were related or unrelated at the time.

Semantically related primes were chosen to be from the same superordinate category as the target word, making them categorically related (e.g. *mouse* and *rat*; *pilot* and *stewardess*), as opposed to words that are merely associatively related (e.g. *dog* and *bone*; *bell* and *tower*). Semantic relatedness was quantified based on two resources, one fully automated and corpus-based, and the other based on published association norms. The automated method was Latent Semantic Analysis (LSA), implemented in on the website http://lsa.colorado.edu/. LSA analyzes the co-occurrence of two terms in a large set of reference documents, and produces a value of cosine similarity reflecting how often the words occur in a similar context (Landauer et al., 1998). The mean estimated cosine similarity for semantically related words in this experiment was 0.45 for nouns in the first position, and 0.39 in the second position, while mean values for phonological primes and unrelated control words ranged from .03 to .08. The norm-based estimates came from the University of Florida free association norms (Nelson et al., 2004), which estimate how frequently various words come to mind given a single prompt word. As this is a directional measure (e.g. dog→cat has a different value than cat→dog), both directions were averaged. The mean association values for semantically related words were 0.20 in the first position and 0.11 in the second, and were uniformly 0 (no reported associations) for phonological primes and unrelated control words. Phonological primes were chosen to match target words in the first two phonemes including the initial vowel (e.g. mouse → mountain, architect → argument), and we verified that no such relationships were present among semantically related pairs or unrelated pairs. Quantitative analysis comparing frequency and length of distractor words and target words in the sentences are presented in supplementary information.

Each participant was presented with a total of 448 trials, organized into 8 runs of MEG recording with 56 trials each, with a run lasting about 7 minutes. Each block contained 8 trials in each experimental condition, plus 8 catch trials, and each sentence frame was used once per run. The first two trials of every run were always catch trials. The order of in which specific items were presented (related or matching unrelated primes first) was counterbalanced across participants in a Latin Squares design, resulting in 8 stimulus lists used across participants, each one being used with two or three participants.

### MEG recording and analysis

Because similar methods have been used in previous studies from our lab (e.g. Kielar et al., 2015; Meltzer et al., 2017), we will only summarize them briefly here. A more detailed description of MEG and MRI processing methods is included in supplementary information.

### MEG recording

MEG signals were recorded with a 151-channel CTF system with axial gradiometers at a sampling rate of 625 Hz. Continuous signals were acquired in 8 runs with brief rest periods between them. Each run comprised 56 trials and lasted approximately 7 minutes. Stimuli in the 9 conditions (8 experimental plus catch trials) were randomly allocated in equal numbers across runs, with 8 different stimulus lists used across subjects. MEG data was coregistered with structural MRI acquired on a 3T scanner at Baycrest, by matching locations of fiducial points. MEG source localization results were transformed into MNI space using a nonlinear warp computed in ANTS (Avants et al., 2011). MEG data were manually screened to identify and exclude trials containing large artifacts related to head or facial movement (e.g. coughs, sneezes, inappropriate speech prior to the recitation cue). Additionally, trials with incorrect sentence repetition responses were excluded. Across subjects, the average number of trials entering analysis was 44-45 for every condition.

### MEG source localization

MEG data was analyzed exclusively in source space, using the beamforming method synthetic aperture magnetometry (SAM, Vrba and Robinson, 2001). For an initial characterization of the time-frequency dynamics induced by our behavioural task, we computed the timecourses of brain electrical activity at a selection of virtual channels in source space, corresponding to the center points of the 90 non-cerebellar regions of the macroanatomical brain parcellation of Tzourio-Mazoyer et al. (2002). For this analysis, beamformer weights were computed based on broadband data (0-100 Hz) spanning the entire time epoch of trials. After conducting the time-frequency and spectral analyses described in the sections below on each virtual channel, we averaged the results across 39 virtual channels that are in left cortical regions. We did not formally consider channels located in the right hemisphere or subcortically.

For whole brain mapping, we compared power in specific frequency bands between different time windows or between different conditions (e.g. sem1rel vs. sem1unr). For these analyses, beamformer weights were computed from data limited to the time-frequency window of interest, resulting in better spatial resolution (Brookes et al, 2008). For each subject, at a regular grid of locations spaced 7 mm apart throughout the brain, we computed the pseudo-T value, which is a normalized measure of the difference in signal power between two time windows (Vrba and Robinson, 2001). More details on the beamforming procedure and spatial normalization of SAM volumes are given in supplementary information. Voxel-wise group statistics on SAM results were computed as a 2-tailed one-sample t-test. To correct for multiple comparisons across the whole cortex, the resulting statistical maps were thresholded on the basis of voxel-wise threshold of p<.01 combined with a cluster-size criterion to achieve a corrected family-wise error rate of p< .05. The necessary cluster size was estimated using a nonparametric permutation procedure, in which the signs of the residuals of the t-test were randomized 10,000 times to estimate the number of expected false positives at a given cluster size and thus generate corrected p-values (Eklund et al., 2016; Cox et al., 2017a). This procedure, implemented in the AFNI program 3dttest++ (calling the program ClustSim implicitly), has been shown to provide accurate estimation of cluster-level family-wise error rates, including at voxel-wise thresholds of p < .01, and is currently the recommended default correction procedure in AFNI (Cox et al., 2017b). For each SAM map presented in the results, we have given the obtained cluster size and estimated cluster-wise p-value in the figure legends. The voxel size of SAM maps in MNI space was 5 mm^3^ isotropic, and clusters were defined for voxels sharing a face or an edge, but not a corner (the default definition in AFNI).

### Time-frequency decomposition

To analyze reactivity to the auditory prime word, single-trial data from virtual channels were decomposed into their time-frequency representation via a Morlet-wavelet transformation implemented in EEGLAB software (Delorme and Makeig, 2004). Two quantities were derived from the wavelet transformation. First, the event-related spectral perturbation (ERSP) quantifies changes in oscillatory power, averaged across trials and normalized with respect to a prestimulus baseline (−300 to −100 ms). ERSP quantifies both power increases (also known as event-related synchronization (ERS) and power decreases, also known as event-related desynchronization (ERD). Second, we computed inter-trial coherence (ITC), a measure of phase synchronization across trials. ITC is useful for distinguishing between non-phase locked oscillatory reactivity, in which case a stimulus induces a change in power but no consistent phase reset of an oscillation, and phase-locked responses, in which case a stimulus evokes a wave of consistent polarity across trials. In cases where a strong ERS and ITC signal are both seen in the same time-frequency window, the signal may be more appropriately characterized in the time domain using the more traditional approach of event-related fields.

### Stationary spectral analysis

To characterize the frequency range of neural reactivity to the task demands, we analyzed power spectra of neural activity at virtual channels during different time periods of the task, including the inter-trial interval (−1.1 to −0.6 seconds prior to visual sentence onset), the initial retention period before onset of the prime word (−0.6 to −0.1 seconds prior to auditory word onset), and the later retention period after the prime word (0.5 to 1.0 seconds after auditory word onset).

For these analyses, we did not do time-frequency decomposition over the entire trial, because the timing of the prime word was variable within the retention period. For each virtual channel and each trial, power spectra were computed using a multitapered Fourier transform. Spectra were averaged across trials at each channel and then averaged across participants. Solely for purposes of display, the “1/f” aperiodic background of the spectra were regressed out (see supplementary information).

### Event-related currents analysis

Analysis of time-domain average signals, typically referred to in MEG as event-related fields (ERFs), was performed in source space over the whole brain on a regular grid of 5mm^3^ voxels, using an adaptation of the event-related SAM (erSAM) technique described by Cheyne (2006, 2007). Because this procedure yields estimates of the underlying electrical currents in the brain that generate the magnetic fields measured by the MEG sensors, we refer to the results as event-related currents (ERCs).

## RESULTS

### Behavioural

The behavioural response in this experiment consisted of the sentence recitation in response to the visual cue at the end of the trial; there was no direct response to the auditory distractor word which occurred during the delay period and was supposed to be ignored. We tested for potential effects of relatedness on (RT, or speech onset latency) and error rate. Error trials included any trials in which the exact sentence was not recited correctly (e.g. a substituted word, an omitted word, a mispronunciation, etc.). Summary statistics are shown in Table 2. We did not analyze catch trials statistically, although we note that RT is much longer on these compared to all the other trial types, likely because of the surprise of seeing the recitation cue earlier than on all other trials, without the intervening distractor.

**TABLE 2:**
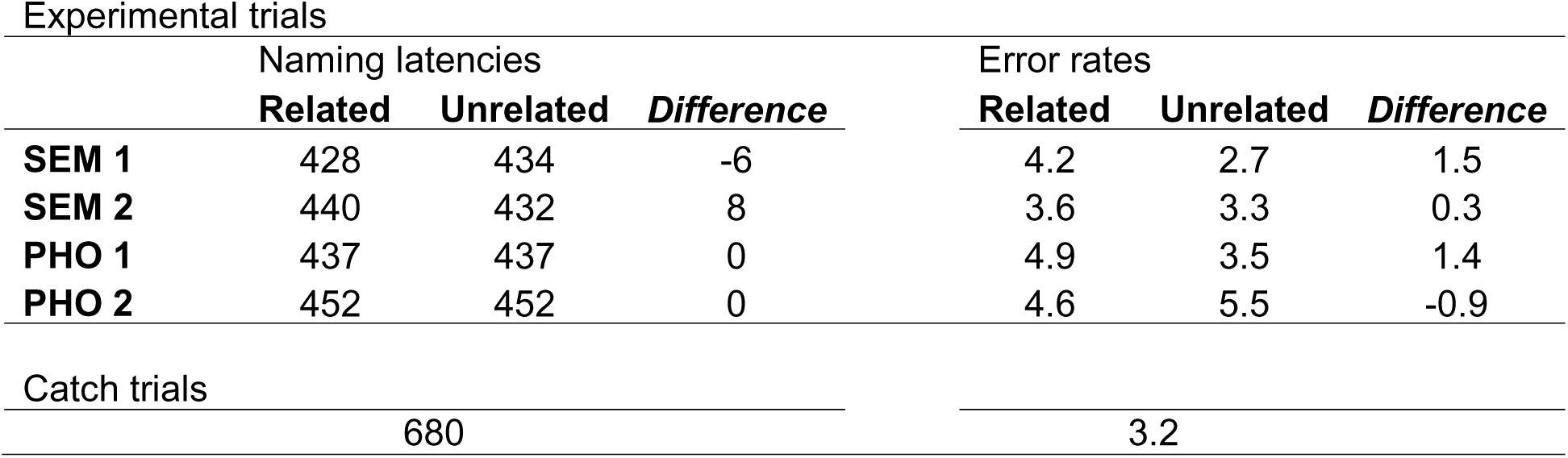
Reaction time (to beginning of utterance) across all conditions. SEM 1,2 = semantically related to first and second noun; PHO 1,2 = phonologically related to first and second noun.

| Experimental trials |  |  |  |  |  |  |
| --- | --- | --- | --- | --- | --- | --- |
|  | Naming latencies |  |  | Error rates |  |  |
|  | **Related** | **Unrelated** | ***Difference*** | **Related** | **Unrelated** | ***Difference*** |
| **SEM 1** | 428 | 434 | -6 | 4.2 | 2.7 | 1.5 |
| **SEM 2** | 440 | 432 | 8 | 3.6 | 3.3 | 0.3 |
| **PHO 1** | 437 | 437 | 0 | 4.9 | 3.5 | 1.4 |
| **PHO 2** | 452 | 452 | 0 | 4.6 | 5.5 | -0.9 |
| Catch trials |  |  |  |  |  |  |
|  | 680 |  |  | 3.2 |  |  |

Error trials were excluded from analysis of RTs. Any RTs deviating from a participant’s or an item’s mean more than three standard deviations were excluded as outliers (39 data points, 0.5% of all trials). RT and error rates were subjected to repeated-measures ANOVA, with relatedness (related/unrelated) and position (noun1 and noun2) as within-subjects factors. Because relatedness type is not meaningful for unrelated words, semantic and phonological relatedness were examined in separate ANOVAs.

No significant main effects or interactions were found for relatedness, either for semantic or phonological stimulus sets. Thus, relatedness of the distractor word does not seem to affect the sentence repetition performance occurring several seconds later.

### Characterization of oscillatory task reactivity

Formal statistical analysis of oscillatory reactivity was carried out on a whole-brain voxel-wise basis across subjects in the range of 8-30 Hz, comparing power between specific conditions. Successful analysis of those differences depends on choosing a time-frequency window that appropriately captures the oscillatory reactivity present in the task. Therefore, we first conducted time-frequency analyses of the response to auditory distractor words at specific virtual channels within the cortex of the left hemisphere. In this analysis, we wished to avoid “double dipping,” a form of circular analysis in neuroimaging in which a researcher might first identify a period of prominent differences and then applying statistical analysis to exactly that period, inflating the statistical significance (Kriegeskorte et al., 2009). To avoid this, we examined the results of these analyses averaged across all eight experimental conditions, avoiding direct comparisons related to the hypotheses. We chose the time-frequency windows in which ERD responses were prominent, without looking at contrasts between the different conditions (e.g. sem1rel vs. sem1unr). Our statistical inferences are limited to the time-frequency windows identified through this procedure.

Time-frequency decomposition of the response to auditory distractor words, averaged across the 39 left cortical channels, is shown in Figure 2A. Auditory words are seen to induce an event-related synchronization (ERS, or power increase), in a low-frequency range from 0-10 Hz, in an early time period from 0 to 0.5 seconds relative to the onset of the word. Additionally, we observe event-related desynchronization (ERD, or power decrease) in a higher frequency range, concentrated between 8-30 Hz as expected. The ERD peaks later than the ERS, and continues from approximately 0.25 seconds all the way to 1.5 seconds, the longest latency analyzed.

**Figure 2.**
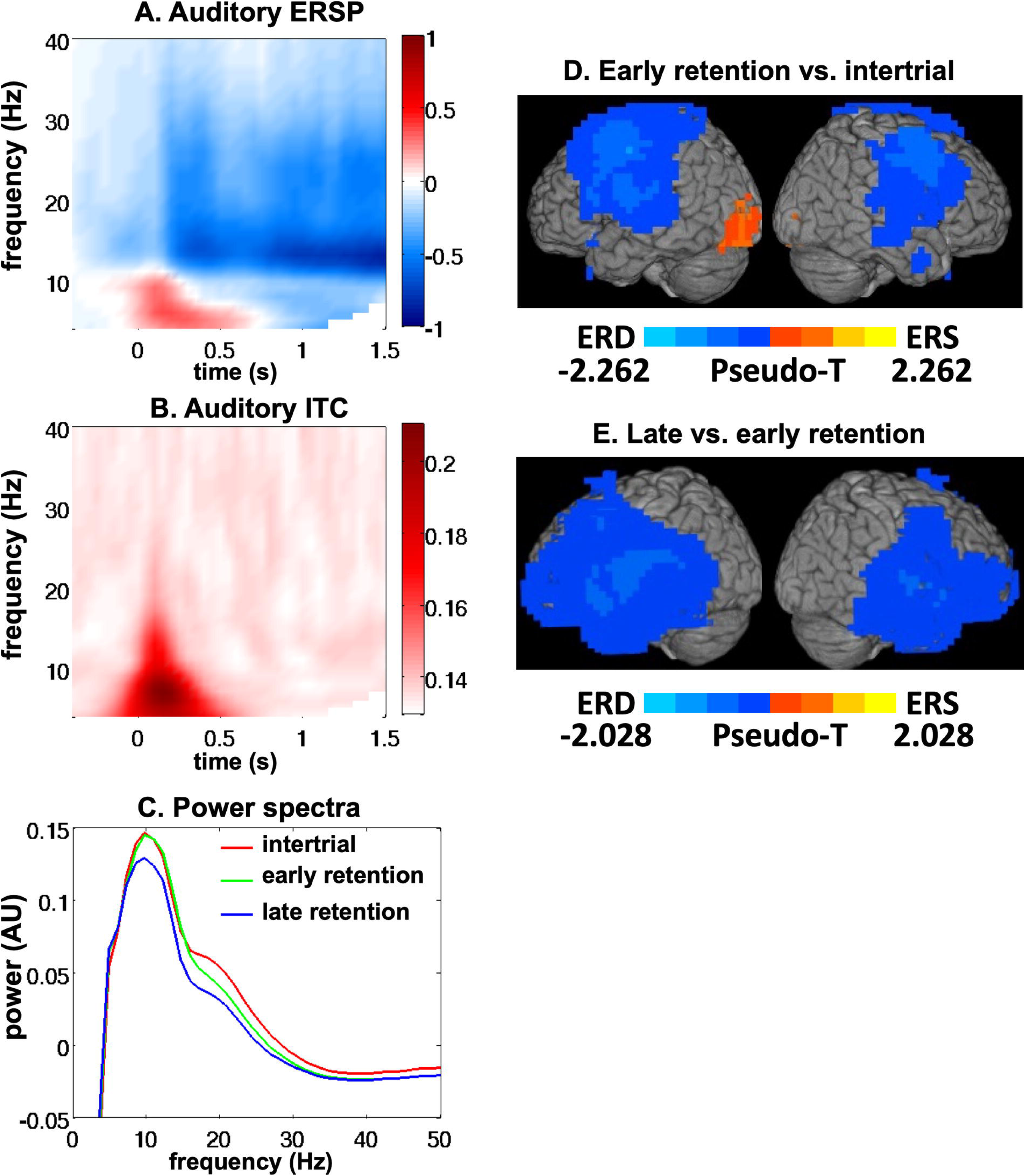
Oscillatory responses irrespective of relatedness. A. Event-related spectral perturbation (ERSP) timelocked to the onset of the auditory distractor word, averaged over all 8 conditions, and averaged over 39 virtual channels in the left hemisphere cortex. B. Inter-trial coherence (ITC), averaged over the same conditions and channels as panel A. C. Power spectra averaged over the 39 cortical channels, extracted from three time periods in the trials – the intertrial interval, the early retention period ((−0.6 to −0.1 s before the onset of the distractor word), and the late retention period (0.5 to 1.0 s after the onset of the distractor word). D. Whole-brain SAM map of 8-30 Hz event-related desynchronization (ERD) in the early retention period vs. the intertrial interval. Voxel-wise threshold p < .01, one significant cluster of size 2916 voxels, cluster p < .01. E. Whole-brain SAM map of 8-30 Hz event-related desynchronization in response to the auditory distractor word, i.e. late retention period vs. early retention period. Voxel-wise threshold p < .01, one significant cluster of size 7815 voxels, cluster p < .01.

To test whether these responses arise from phase-locked signals as opposed to simply induced changes in power, we examined ITC for the same signals, shown in Figure 2B. A strong phase-locked signal is observed in low frequencies, mainly between 0-10 Hz but ranging up to 20 Hz at some time-points. This signal dissipates by 0.5 seconds. Thus, the early 0-10 Hz ERS visible in Figure 2A appears to be phase-locked, and may appropriately be examined in the time domain using analysis of event-related currents. In contrast, the 8-30 Hz ERD has no counterpart in the ITC map, suggesting that it is a purely oscillatory phenomenon.

In addition to the responses to the auditory distractor word, we were also interested in power differences between the memory delay period (both before and after the auditory word) and the intertrial interval. Differences between these time periods should reflect brain activity related to holding the sentence in working memory, and processing the auditory word (although it should be ignored). The timing of this activity is constrained by the task design, but it is uncertain whether the frequency ranges involved are identical to those responding to the auditory word. To characterize the responsive frequency range, we conducted power spectral analysis on activity reconstructed at the same 39 channels during different periods of the task (see methods). The results are shown in Figure 2C. As expected, oscillatory power is reduced during the initial retention period compared to the inter-trial baseline, and then it is reduced further after presentation of the auditory prime word. Although these differences extend into higher frequency ranges, their magnitude is maximal in the range of 8-30 Hz, again as expected. Therefore, we conducted whole-brain statistical analyses of task reactivity in the range of 8-30 Hz.

### Whole brain task responses

Using whole-brain SAM to compare oscillatory power between different time windows, we compared 8-30 Hz power during the early retention period (−0.6 to −0.1 seconds prior to auditory word onset) with the pre-stimulus baseline period (−1.1 to −0.6 seconds prior to visual sentence onset). The resulting activation map is shown in Figure 2D.

An extensive bilateral pattern of ERD was seen, with maximal reactivity in the frontal lobe, especially in the precentral gyrus. Although the pattern was bilateral, it was stronger on the left. Positive signal change, i.e. 8-30 Hz ERS (power increase), was observed in the left occipital lobe, extending into the cerebellum. This occipital ERS effect may be a “rebound” of power after strong ERD induced by the visual word stimuli, and will not be considered further.

We next compared activity following the prime word with that occurring before the prime word (0.5-1.0 s post-onset vs. −0.6 to −0.1 s pre-onset). Again, a strong pattern of bilateral ERD was induced (Figure 2E). The extent of ERD covered most of the temporal lobes in both hemispheres, and extended into the frontal lobes, especially on the left. In general, ERD induced by the prime word was slightly stronger in the left hemisphere, and maximal in the left temporal lobe. No positive signal changes (ERS) were observed for this contrast.

### 8-30 Hz ERD: Effects of phonological and semantic relatedness

Based on the analyses presented above, we elected to analyze differences between conditions in a window of 8-30 Hz and 500-1000 ms after the onset of the auditory distractor word. This time-frequency window was chosen based on the presence of strong ERD in the analyses described above, without regard to differences between conditions. Effects of phonological and semantic relatedness were analyzed separately. For each subject, we generated four whole-brain maps of pseudo-T values reflecting the difference in power between two conditions. These maps are referred to as:

1. Pho1 = pho1rel – pho1unr
2. Pho2 = pho2rel – pho2-unr
3. Sem1 = sem1rel – sem1unr
4. Sem2 = sem2rel – sem2unr

Thus, for each subject there were maps of the difference in power after phonologically related and unrelated words in noun position 1 and 2, and the same for semantic relatedness. For each kind of relatedness, two planned contrasts were conducted.

Main effect of phonological relatedness = (Pho1 + Pho2) / 2
Phonological position effect (interaction) = (Pho1 – Pho2)
Main effect of semantic relatedness = (Sem 1 + Sem2) / 2
Semantic position effect (interaction) = (Sem1 – Sem2)

These two planned contrasts were used instead of a full factorial ANOVA, as we were not interested in the differences between the unrelated conditions (i.e. pho1unrel vs. pho2unrel). For statistical analyses across participants, we first computed the planned contrast maps for each participants using the formulas presented above, and then subjected the results to a one-sample t-test across subjects, corrected for multiple comparisons using cluster-based permutation tests (see methods).

The main effect of phonological relatedness is shown in Figure 3A. Compared to unrelated words, auditory distractor words that were phonologically related to a noun in the memorized sentence induced greater 8-30 Hz ERD in extensive portions of the left hemisphere cortex, and to a much lesser extent in the right hemisphere. Activation levels are highest throughout the left inferior frontal gyrus, including Brodmann Areas (BA) 44, 45, and 47. From there activation extends posteriorly and superiorly into the Rolandic Operculum, precentral and postcentral gyri, superior and middle temporal gyri, supramarginal gyrus, inferior parietal lobe, angular gyrus, superior parietal lobe, and precuneus. Activity is also seen in the left middle frontal gyrus, middle cingulate, and supplementary motor areas. In the right hemisphere, activity is largely confined to the middle frontal gyrus, extending inferiorly to portions of the right inferior frontal gyrus, BA 44 and 45. Only negative values are seen; there were no regions exhibiting higher 8-30 Hz power in response to phonologically related words.

**Figure 3.**
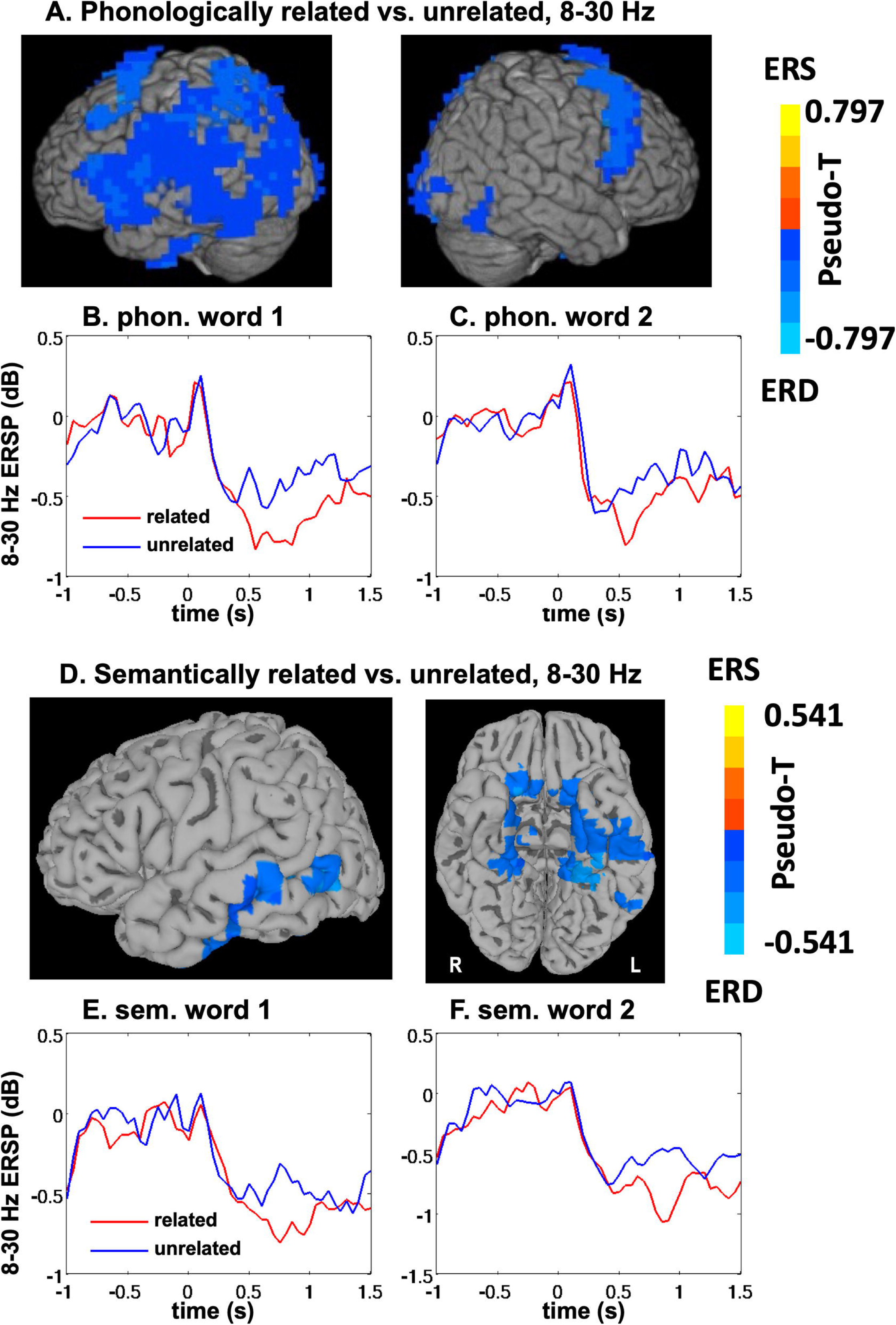
Effects of relatedness on 8-30 Hz ERD. A. Whole-brain SAM map of 8-30 Hz oscillatory power, 0.5-1.0 seconds after auditory distractor word onset, for phonologically related vs. unrelated distractors. Effects are averaged across words related to the first and second nouns in the sentence. Voxel-wise threshold p < .01, one significant cluster of size 3998 voxels, cluster p < .01. B. Timecourse of 8-30 Hz power at the virtual channel closest to the peak of the phonological relatedness effect, timelocked to the onset of the auditory distractor word, for distractors phonologically related to the first noun of the visual sentence. C. Timecourse for distractors phonologically related to the second noun of the visual sentence. D. Whole-brain SAM map of 8-30 Hz oscillatory power, 0.5-1.0 seconds after auditory distractor word onset, for semantically related vs. unrelated distractors. Effects are averaged across words related to the first and second nouns in the sentence. Because much of the effect is in more medial portions of the temporal lobe, significant voxels are overlaid on an inflated pial surface shown from left and ventral views. Voxel-wise threshold p < .01, one significant cluster of size 707 voxels, cluster p < .02. E. Timecourse of 8-30 Hz power at the virtual channel closest to the peak of the semantic relatedness effect, timelocked to the onset of the auditory distractor word, for distractors semantically related to the first noun of the visual sentence. F. Timecourse for distractors semantically related to the second noun of the visual sentence.

Although there was a strong main effect of phonological relatedness, we did not detect a significant interaction effect. The contrast (Pho1 – Pho2) returned no significant clusters (data not shown). Thus, the effect of phonological relatedness appears to be roughly equivalent whether the distractor word is related to an early or late noun in the memorized sentence, within the statistical power achieved in the present study (see supplementary information for sensitivity power analysis).

To illustrate the timecourse of this effect, we plotted ERSP in the 8-30 Hz range from the virtual channel that was closest to the peak voxel of the whole-brain map. The peak voxel was located in left occipitotemporal cortex at [MNI −33 L, −71 P, 10 S], and the virtual channel was that corresponding to the left superior temporal gyrus in the AAL atlas (region 81). Timecourses for ERSP in response to related and unrelated words are shown in Figure 3B for the first noun, and 3C for the second noun.

Consistent with the pattern seen in the averaged ERSP over all left cortical channels (shown in Figure 2A), auditory distractor words induce a brief increase in power (ERS), followed by a prolonged decrease (ERD) that lasts at least as long as 1.5 s, the longest latency analyzed. The ERD is stronger for related words. The difference between related and unrelated words subsides after about 1 s, although power for both conditions remains suppressed relative to the period before word onset. The response patterns for words related to the first and second noun are very similar, which is consistent with the lack of an interaction effect in the whole-brain statistical analysis.

The main effect of semantic relatedness is shown in Figure 3D. Compared to unrelated words, auditory distractor words that were semantically related to a noun in the memorized sentence induced greater 8-30 Hz ERD. Unlike the phonological effect however, the semantic effect was rather limited in extent, concentrated mainly in the left temporal lobe, specifically in the middle temporal gyrus, but extending along the ventral surface of the brain into the fusiform gyrus and into medial portions of the right temporal lobe. Due to the spatial imprecision of MEG source analysis, it is difficult to know if the right hemisphere involvement actually reflects neural activity in the right temporal lobe, or instead represents “bleeding” of sources in the left hemisphere. As much of this activity is located in more medial regions, we display it on a slightly inflated pial surface to make it more visible. As with the phonological effects, these differences were exclusively negative: no regions showed greater 8-30 Hz power in response to semantically related words.

Also similar to the phonological effects, there was no interaction effect with noun position for semantic relatedness. The contrast (Sem1 – Sem2) returned no significant clusters (data not shown). Thus, the effect of semantic relatedness appears to be roughly equivalent whether the distractor word is related to an early or late noun in the memorized sentence, given the statistical power achieved in this study.

To illustrate the timecourse of this effect, we plotted ERSP in the 8-30 Hz range from the virtual channel that was closest to the peak voxel of the whole-brain map. The peak voxel was located in the left middle temporal gyrus at [MNI −43 L, −61 P, 0 S], and the virtual channel was that corresponding to the left middle temporal gyrus in the AAL atlas (region 85). Timecourses for ERSP in response to related and unrelated words are shown in Figure 3E for the first noun, and 3F for the second noun. In response to the auditory distractor word, 8-30 Hz power is suppressed, and suppressed more strongly for semantically related words, especially in the 0.5-1.0 second range that the statistical analysis was based on.

### Event-related currents: Effects of phonological relatedness

Prior electrophysiological studies of semantic and phonological priming for auditory words have typically shown effects ranging from 300 to 1000ms after word onset (e.g. Bouaffre and Faita-Ainseba, 2007; Jescheniak et al., 2002; Phillips et al., 2006; Von Holzen and Mani, 2014). These effects fall into the traditional time windows of the ERP effects known as N400 (approximately 350-550 ms), which is typically suppressed by semantic relatedness and is thought to be generated primarily in the left temporal lobe (Lau et al., 2008), and the P600 (approximately 600-1000 ms) which has been shown to be modulated in various ways by linguistic manipulations, and has a more uncertain localization. However, auditory priming effects on ERPs typically take the form of signal shifts across a wide time range, rather than changing the amplitude of a discrete peak. This may be due to less precise time-locking of neural activity compared to visual presentation, as the point at which a word is recognized (the “uniqueness point”) varies greatly between words (Radeau et al., 1989). For an initial analysis, we tested for differences (main effects and interactions, as described above for the ERD analysis) between related and unrelated words in two time windows, an earlier one from 0.4 to 0.6 s, and a later one from 0.6 to 0.9 s. However, the results of these two analyses were nearly identical, identifying essentially the same cluster of significant activity, so we replaced them with a single analysis covering the entire time window from 0.4 to 0.9 s.

One significant cluster was found for a main effect of phonological relatedness, centered in the left precentral gyrus, shown in Figure 4A. This took the form of a larger response for related vs. unrelated words. To illustrate the timecourse of event-related currents driving this effect, we extracted the averaged signals from a spherical ROI of 8 mm centered on the peak voxel of the cluster, located at [MNI −40 L, −16 P, 42 S]. The peak location was used instead of the closest atlas-based virtual channel, because the ERCs were already computed on the whole-brain level, unlike time-frequency decompositions. The timecourses for ERCs in response to words phonologically related to the first and second words of the memorized sentence, along with the responses to the unrelated control words, are shown in Figure 4B and 4C, respectively. Stronger responses are seen to related words for both positions. Notably, the difference seems to be more protracted in duration for words related to word 1 compared to word 2. However, no significant interaction effect for relatedness by position was found in the time window that we investigated.

**Figure 4.**
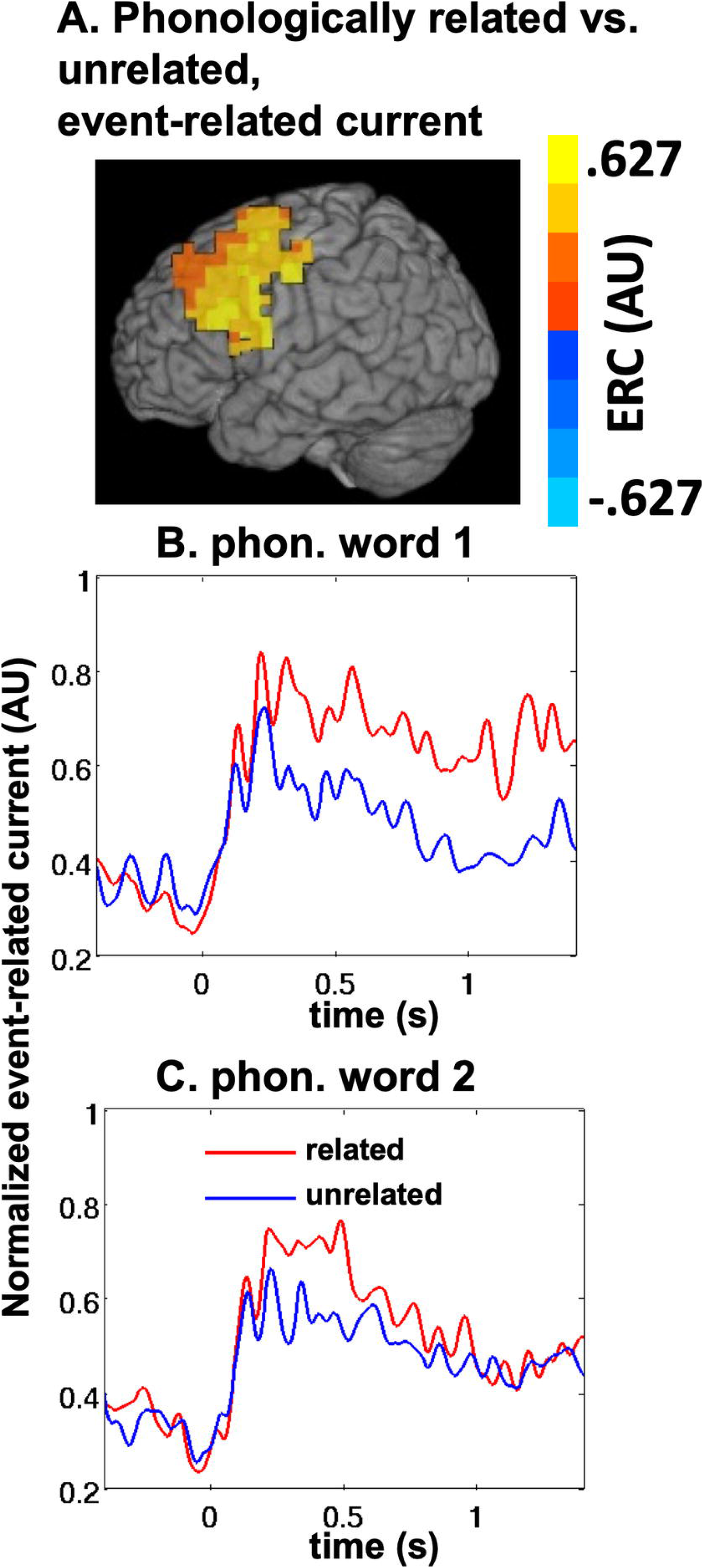
Effects of relatedness on event-related currents. A. Whole-brain SAM map of time-domain event-related currents, 0.4 to 0.9 s after auditory distractor word onset, for phonologically related vs. unrelated distractors. Effects are averaged across words related to the first and second nouns in the sentence. Voxel-wise threshold p < .01, one significant cluster of size 304 voxels, cluster p < .01. B. Timecourse of event-related currents at a virtual channel near the peak of the phonological relatedness effect, timelocked to the onset of the auditory distractor word, for distractors related to the first noun of the visual sentence. C. Timecourse for distractors related to the second noun of the visual sentence.

For semantic relatedness, we did not find any significant main effect or interaction in the analysis of event-related currents (data not shown).

## DISCUSSION

As outlined in the introduction, the present study was designed to address three specific questions about advanced planning of utterances under conditions in which the entire utterance is held in short-term memory awaiting the cue to speak.

### 1) Spatial distribution of semantic and phonological representations

Our preliminary analyses of oscillatory reactivity in this paradigm, agnostic of relatedness conditions, indicated that 8-30 Hz oscillatory power in a wide range of cortical areas was suppressed during maintenance of sentences in short-term memory, consistent with previous MEG studies employing sentence maintenance (e.g. Meltzer et al., 2017; Meltzer and Braun, 2011). Additionally, we observed that presentation of the auditory prime word induced a further decrease of power in the same range. We examined differences in the magnitude of this auditory event-related desynchronization as a function of semantic and phonological relatedness.

We hypothesized that semantic and phonological relatedness would exert distinct effects in brain networks that specialize in the maintenance of these two kinds of information. Brain areas involved in STM for verbal repetition have been reviewed by Majerus (2013), who classifies them into three distinct networks. A “ventral language pathway” includes left middle temporal (BA 21/38) and ventral left inferior frontal (BA 45) areas, and is thought to implement maintenance of semantic item information. A “dorsal language pathway,” including left superior temporal (BA 22) and left ventral inferior frontal (BA 44) areas, underlies maintenance of phonological item information, including both words and nonwords. To these two language pathways, Majerus adds a bilateral “fronto-parietal network,” (BA 9, 6, 40, 7) involved in maintenance of serial order information and allocation of attentional control to the task, particularly for repetition of multiple words and nonwords. The combination of these three pathways largely overlaps with the dual-stream model of language networks (not specific to repetition) proposed by Hickok and Poeppel (2004, 2007), and shares with it the specialization of the ventral pathway for lexical semantics, with more dorsal areas involved in processing phonological representations and transforming them into articulatory codes. Thus, we predicted that semantic relatedness would modulate auditory responses in ventral areas, and phonological relatedness would modulate them in dorsal areas.

Our results were broadly consistent with this prediction, but did not indicate a full double dissociation. For *semantic relatedness*, as expected, we observed modulation only within the ventral network.

Semantic relatedness induced an enhanced ERD response in a single cluster confined to the temporal lobe, concentrated mainly within the left middle temporal gyrus (MTG). This region has been implicated in several prior neuroimaging studies as being critical to semantic aspects of word production. In a series of meta-analytic reviews exploring the spatial and temporal characteristics of neural activity involved in word production (Indefrey and Levelt, 2000; 2004), the left MTG was posited to be a critical locus of conceptual/semantic retrieval based on findings that, out of several areas reliably activated in neuroimaging studies of picture naming and word generation tasks, the MTG alone is not generally recruited in studies of reading written words, consistent with it being involved in generation of a semantic representation of a specific word (a “lemma”) from conceptual rather than perceptual information. In a later review (Indefrey, 2011), this “negative evidence” is supplemented by positive findings pointing to a specific semantic role of the left MTG in word production. These include enhancement of fMRI activation for semantically related distractors in picture-word-interference tasks (de Zubicaray et al., 2002; 2006), enhanced MEG responses (presumably reflecting semantic competition) during picture naming when pictures were presented in blocks of categorically related items (Maess et al., 2002), and the identification of the left anterior-to-middle temporal gyrus in a voxel-based lesion-symptom mapping study of picture naming by stroke patients as a critical region, in which a lesion predicts the production of semantic errors in the absence of non-verbal conceptual deficits (Schwartz et al., 2009).

The present finding of selectively enhanced responses in the left MTG to words that are semantically related to the memorized sentence suggests that this area plays a role not only in the *retrieval* of conceptual information in word production, but also in semantic aspects of *maintenance* for planned utterances held in short-term memory. Although maintenance of a short sentence (5 words in the present study) is likely to rely mainly on rote rehearsal, i.e. phonological short-term memory, the enhanced response suggests that ongoing neural activity related to the sentence’s meaning is also present during the delay period within the left MTG, and the perception of a related word interacts with this ongoing activity. This is true even though the participant is free to ignore the auditory word completely, as it has no relevance to correct performance of the experimental task.

Another possible interpretation of the enhanced ERD in response to semantically related distractors is that the involved brain regions are simply involved in resolution of interference from the distractor word, and not necessarily in *maintenance* of the semantic information. This distinction relates to a long-running controversy on the role of the posterior MTG (pMTG) in semantic processing. Several studies examining lesion evidence and effects of “virtual lesions” with transcranial magnetic stimulation (TMS) have suggested a role for this region in semantic control processes, such as selecting the less common meaning of a word (e.g. *bank* referring to a river bank instead of a financial institution) or selecting a weak semantic associate (e.g. *salt* and *grain*) vs. a strong associate (e.g. *salt* and *pepper*), (Jefferies et al., 2006; Whitney et al., 2011; Hoffman et al., 2012). This special role in executive control over semantics contrasts with more general cognitive control functions subserved by the multiple demand network (Davey et al., 2016) and with specialization for storage of broad semantic representations independent of control demands, which is associated with the anterior temporal lobe (Lambon Ralph et al., 2017). However, other studies have highlighted other roles for the pMTG, including processing of compositional relations among multiple items (e.g. compound words, or nouns vs. verbs) and temporal integration of meanings in continuous language comprehension (Matchin et al., 2019; Bedny et al., 2014; Guediche et al., 2013).

Our identification of this region as involved in resolution of semantic interference during short-term memory for sentences adds to this picture of the pMTG as a region essential for processing of semantics in a temporally extended context. It also converges with evidence from a previous, separate MEG study from our lab that also examined sentence repetition, using longer sentences without any distractors (Meltzer et al., 2017). In that study, we attempted to identify brain regions involved in maintenance of semantic information by contrasting activity during the delay period while participants were maintaining highly imageable (i.e. concrete) sentences vs. abstract sentences. We reasoned that the more imageable sentences were likely to benefit from semantic support and mental imagery in their maintenance, whereas the abstract sentences would be more dependent on rote phonological rehearsal, a prediction born out by a companion behavioural study (Meltzer et al., 2016). We found enhanced 8-30 Hz ERD in the posterior temporal lobe, overlapping heavily with the cluster of semantic relatedness in the present study, although in that case the activation was bilateral. Together, these two MEG studies converge on the conclusion that the left posterior MTG plays a key role in the maintenance of semantic information supporting accurate sentence repetition.

In contrast to the relatively circumscribed area of enhanced response to semantically related words, *phonological relatedness* exerted strongly enhanced 8-30 Hz ERD in a widespread network of left-hemisphere cortical regions, with additional involvement of the right hemisphere precentral gyrus (comprising primary motor and premotor cortices). The activated regions include portions of all three networks identified by Majerus, including the right precentral gyrus for the bilateral fronto-parietal network. This much larger pattern of activation may be related to the nature of the task, in that participants are most likely engaging in rote rehearsal of the sentence throughout the delay period, and thus recruiting entire networks that are typically activated in studies requiring the use of phonological short-term memory to maintain verbal material through a delay period. Our task, with five-word sentences as repetition material, could be expected to engage all of these regions, as participants must maintain both item identity and sequence information to correctly reproduce the sentence.

It should be noted that middle temporal regions that are modulated by semantic relatedness are also modulated to a similar degree by phonological relatedness; direct statistical contrasts showed that there were no regions in which the semantic effect was significantly larger than the phonological effect. Thus, the present results do not reach the level of double dissociation. Ventral regions are modulated by semantic relatedness as expected, but they are also equally modulated by phonological relatedness. One possible explanation of this pattern is that words that are phonologically related to words in the maintained sentence may also lead to activation of the target word as well as semantically related associates of it, giving rise to a semantic interference effect in addition to a phonological interference effect, but the reverse is not true: semantic distractors do not necessarily induce phonological interference. However, this account is speculative and would need to be confirmed through further experiments. Here again, it is useful to compare the present results with our previous MEG study of sentence repetition, Meltzer et al., (2017). In that study, we attempted to map regions associated with phonological STM by comparing activity during maintenance of arbitrary word lists vs. structured sentences. The resulting activation map (see Figure 2D-E of that paper) was largely similar to that seen in the present work for phonological relatedness in the left hemisphere (Figure 3A), but with notably *more* involvement of the middle and inferior temporal lobe in the present study, suggesting a possible role of these regions for interference resolution beyond simple maintenance of verbal information. Also, in the previous study, maintenance of word lists induced massively increased ERD in the right hemisphere, whereas in the present study, resolution of phonological interference induced relatively limited right hemisphere ERD, mainly in the precentral gyrus. The increased right hemisphere activation seen in the previous study may be attributable to greatly increased demands on pSTM when maintaining word lists with no semantic support, compared to the more subtle manipulation of resolving interference in the present study.

In addition to the modulations of 8-30 Hz ERD induced by both kinds of relatedness, we also observed a significant modulation of the time-domain event-related currents only in response to phonological relatedness (see supplementary information for discussion of statistical power limitations regarding the null finding for semantic relatedness). Larger responses for phonologically related distracters were observed in a relatively circumscribed cluster centered on the left precentral gyrus. This area is thought to be involved in motor planning including for speech (Indefrey and Levelt, 2004). Thus, a larger response in this area is consistent with the idea that phonological distractors induce interference in the speech planning system, requiring areas involved in articulatory planning to process auditory input more intensively in order to resolve the interference.

### 2) Temporal scope of advanced planning for semantic and phonological representations

As described in the introduction, some studies have suggested that the temporal scope of advanced planning for speech is larger for semantic than for phonological information. Given this background, we hypothesized that semantic relatedness effects would be equally strong for early and late words in the sentence, but that phonological relatedness would primarily exert an effect for early words. This prediction was not confirmed. Instead, we found robust relatedness effects for both early and late words in the sentence, and no significant difference in the magnitude of the relatedness effects across word positions, within the power achieved in the present study (see supplementary information for a discussion of effect sizes and statistical power within this experiment).

In interpreting this finding, we must point out that our repetition task differed in several respects from the speech planning studies that have shown differences in temporal scope. These studies are largely based on observations of spontaneous speech errors in natural speech, or on tightly controlled chronometric picture naming studies. In both of those forms of speech, the intended utterance is not strictly determined by the experimenter, but rather by the participant. Thus the speech act includes retrieval of the intended words, rather than simply maintaining them in memory. In our experiment, the complete utterances were already encoded verbatim into short-term memory before participants heard the distractor words. Thus, one could argue that “planning” of the utterance is already complete, and that the full form is simply being maintained until the repetition act, and no further “advance planning” is occurring at this stage.

Another consideration is the fact that in delayed sentence repetition, the internal mental generation of the words is somewhat unconstrained. During the delay period, participants are most likely engaging in rote rehearsal of the sentence, and given variability in the speed of rehearsal, the exact word that they are thinking of in their covert speech will vary at the point when the distractor is presented. Any potential differences in advance planning might be washed out by this variability. It is therefore possible that a modified task paradigm that more tightly constrains the timing of speech planning might be more effective at showing differences in the scope of advanced planning of speech on the semantic and phonological levels. For example, participants could be cued to produce one word at a time, and the distractor word could be presented (near-)simultaneously with the cue to produce the planned utterance, at different positions within the sentence. However, this technique would risk introducing speech-related artifact at the same time as the participant is processing the distractor.

### 3) Directionality of effect

In all regions, related words induced a stronger ERD (power reduced to lower levels) compared to unrelated words. Interestingly, these differences were observed in the absence of any behavioural effects – the presence of related vs. unrelated distractors during the memory delay did not affect participants’ performance in reciting the sentence. Most likely, this is because any facilitation or interference produced by the distractor words was fully resolved before the participants were cued to recite the sentence. This is consistent with behavioural findings from PWI tasks, which typically report maximal semantic interference when a distractor word precedes the target picture by a few hundred milliseconds, and maximal phonological facilitation when the distractor and target are presented simultaneously. Both of these effects disappear with longer delays before target presentation, testing with stimulus-onset asynchronies up to +300 ms (see Abel et al., 2009 for review). Thus, although the behavioural output in this study occurs long after the interference has faded, the electrophysiological responses to the distractor words are measured instantaneously upon distractor presentation, and can show evidence of facilitation or interference without overt behavioural effects.

As we observed only increased ERD in response to related words compared to unrelated words, prior knowledge about electrophysiological responses suggests that both types of distractors likely introduced interference. Typically, greater ERD in the alpha and beta bands corresponds to an increased BOLD response in fMRI, and to a higher neural firing rate (Singh et al., 2012) suggesting more intensive processing of a stimulus. Generally, this is associated with more difficult conditions, e.g. conditions creating more interference and leading to worse behavioural performance. Semantic and phonological distractors are of great interest in psycholinguistics because, when presented at appropriate time intervals relative to a picture in PWI studies, they can induce opposite effects: semantic interference and phonological facilitation (Abel et al., 2009; Schriefers et al., 1990; Wei et al., 2022; Jescheniak and Schriefers, 2001). Thus, in the present study we expected that conditions of interference would lead to greater ERD.

The finding that both kinds of distractors appear to generate interference in the present study, rather than opposite effects as seen in PWI studies, stems from the difference in task demands. In PWI studies, the participant is asked to retrieve a lexical form from a purely semantic cue, i.e. a picture. While semantic distractors activate representations of close competitors, creating interference, phonological distractors activate phonemic representations that overlap with the target word, making it easier for the participant to retrieve that target word. In the present study, however, lexical retrieval is not required from the participants. During the memory delay, they had just read the sentence and were only required to repeat it verbatim. Thus, all of the words in the sentence had effectively *already* been retrieved, and were all active in short-term memory as the participants rehearsed the sentence. Sentence repetition relies on both phonological short-term memory (pSTM) and semantic, or conceptual, short-term memory (cSTM). Although pSTM may be sufficient for maintenance of short utterances like the ones used in this study, it is well-known that semantic resources can supplement pSTM, leading to phenomena like the sentence superiority effect, in which participants can remember much longer sentences compared to arbitrary word lists, as the memory for the overall meaning of the sentence allows for the surface form of the sentence to be “redintegrated” into pSTM (Baddeley et al., 2009; Allen et al., 2018). Another form of evidence for cSTM is that patients with damage to systems supporting pSTM often lose the ability to repeat sentences verbatim, but can generate recognizable paraphrases maintaining the overall meaning (Baldo et al., 2008).

In the present study, it appears that related distractor words create increased interference specifically in anatomical regions involved in maintenance of phonological or semantic information. Given that the sentences are short and can probably be maintained easily through pSTM alone, it is unsurprising that phonological distractors appeared to have a much stronger effect, enhancing 8-30 Hz ERD throughout the language network and in additional areas, predominantly but not exclusively in the left hemisphere. In contrast, the effects of semantic distractors were more limited in extent, confined entirely to the temporal lobe, also with a left-hemisphere preponderance but extending to medial portions of the right temporal lobe. This finding is consistent with the “dual-stream” model of language representation, in which semantic computations take place bilaterally throughout the temporal lobe, proceeding anteriorly and becoming increasingly multi-modal and abstract, culminating in high-level semantic representations housed in the anterior temporal lobe, and making contacting with other aspects of language through projections to the ventral frontal lobe (Hickok and Poeppel, 2007; Lambon Ralph et al., 2017). Maintenance of phonological information, in contrast, is thought to arise from activation of more dorsally located and left-lateralized portions of the temporal, parietal, and frontal lobes, engaging regions that are also involved in overt speech production (Hickok and Poeppel, 2007).

## Limitations

Some specific limitations of the present study must be acknowledged. First, the sample size of 20, while fairly typical for language neuroimaging studies, may have been inadequate to capture some subtler differences in neurophysiological activity related to the experimental conditions. One example is that of ERCs for semantic distractors vs. unrelated words, for which no significant clusters were detected. Another is position effects, in which the effect of phonological relatedness might be more prominent for the first noun in the sentence compared to the second, reflecting a limited scope of advance planning for speech. Hints of such an effect can be seen in Figure 4B-C, where the effect of phonological relatedness on ERC signals in the left frontal lobe appears to be longer lasting for the first word compared to the second. This difference was noted visually, but did not reach statistical significance. It is possible that a study with a larger sample size could clarify whether this effect is real. More information about the effect sizes observed within the study and the achieved statistical power is presented in supplementary information.

Another limitation relates to the fundamental nature of the MEG signal – it is sensitive to some kinds of neural activity but not all. It is exceptionally good at detecting differential ERD in the alpha and beta bands, for example, as seen in this study, but relatively poor at detecting increased power in the high gamma band (e.g. 60-100 Hz), an effect that seems to be very robustly detected for language paradigms in human intracranial recordings (Crone et al., 2011). Thus, it is possible that an alternative recording modality such as intracranial EEG could reveal additional effects not seen in the present study.

## Conclusion

Both semantic and phonological distractors produce short-lived interference with word representations during rehearsal of short sentences. These effects are differentially localized, to bilateral temporal lobes for semantic distractors, and a broader left-lateralized network for phonological distractors. However, the distribution of effects, while consistent with a dual-stream view of sentence maintenance, does not demonstrate a full double dissociation, with the temporal areas showing both semantic and phonological effects. This study did not demonstrate differences in temporal scope of semantic and phonological planning, but this may be attributable to the unconstrained nature of rehearsal during the delay period. Future work with more constrained sentence planning tasks may reveal differences in the scope of semantic and phonological aspects of word retrieval during spontaneous or cued speech.

## Data Availability Statement

Materials for this study are available at https://osf.io/cafvk/. The upload includes the raw BIDS format MEG and defaced structural MRI data for all participants, a list of trigger codes for the experimental conditions as embedded in the MEG files, and the full list of experimental stimuli.

