## supplementary information for "Oscillatory brain activity reflects semantic and phonological activation during sentence planning"

**Comparison of distractor words and target words.**

The primary comparisons of interest in the present study concerned distractor words that are semantically or phonologically related to target words in the sentences that participants had to maintain in short-term memory and then recite on cue. Because there were numerous constraints to choosing such words, we designed the study such that the related and unrelated words *were the same words*, counterbalanced across sentences so that they would be related in one sentence and unrelated in a different one. Thus, all comparisons between related and unrelated words involved the same words in different contexts. Nonetheless, a reviewer pointed out that the distractor words may or may not be matched in lexical properties to the target words in the sentences. To assess this, we quantified three key properties for each distractor word, as well as the two target words (the subject and object nouns) in each sentence. We estimated word frequency from the SUBTLEX database (Brysbaert and New, 2009), length in syllables from the CMU pronunciation dictionary (Weide, 1998) implemented in the python package NLTK (Bird et al., 2009), and length in characters (measured directly). Using the 48 sentence frames, we compared the subject and object target words with the following distractors: pho1rel, pho1unr, sem1rel, sem2rel, pho2rel, pho2unr, sem2rel, sem2unr, using a series of 8 paired t-tests.

For word frequency, we found that pho2rel (p=.005) and pho2unr (p=.008) were less frequent than the subject, and pho1unr (p=.036), pho2rel (p=.036), and pho2unr (p=.032) were less frequent than the object. Thus, the phonological distractor words tended to be less frequent than the target words in the sentences. This is unsurprising, given that choosing words with the same phonological onsets as target words in sentences is highly constraining, and some of the words chosen are somewhat obscure (e.g. *chieftain* for *cheese*, *howler* for *house*). Given these constraints, our decision to use counterbalancing across sentences with the same distractor words makes sense. There were no significant differences between semantic distractor words and target words for word frequency.

For length in syllables, the only differences were that sem1rel (p=.004) and sem1unr (p=.012) were shorter than the corresponding subject words. No differences were observed compared to the object words, or for phonological distractors. This may relate to a tendency among the experimenters to prefer monosyllabic words when thinking of potential semantic matches for target words. For length in characters, there were no significant differences for any of the comparisons.

**MEG recording**

MEG signals were recorded with a 151-channel whole-head system with axial gradiometers (CTF, Coquitlam, BC, Canada). MEG was recorded continuously at a sampling rate of 625 Hz, and acquired with online synthetic 3rd-order gradient noise reduction (Vrba and Robinson, 2001). Continuous signals were epoched based on specific timepoints of the trial, described in the results section. Head position with respect to the MEG helmet was monitored using three coils placed at anatomical landmarks of the head (nasion, left and right pre-auricular points). The head position was measured before and after each run and averaged across runs for source analysis. The averaged maximal amount of RMS motion between any two runs for any coil was 7.2 mm.

**MRI acquisition and processing**

Each subject underwent a one-hour structural MRI session on a 3T scanner (Siemens TIM Trio) located at Baycrest. The one-hour sessions included several clinically oriented scans collected for use as a healthy control group for other studies, which are not discussed here. A high-resolution T1-weighted anatomical scan was used to construct a head model for MEG source modeling (MPRAGE, 1 mm isotropic voxels). MR-visible markers were placed at the fiducial points for accurate registration, aided by digital photographs from the MEG session. MRI was typically acquired 1–14 days after the MEG session. To construct head models for MEG analysis, the locations of the fiducial points were marked manually in AFNI software (Cox, 1996), and the T1-weighted MRI was spatially transformed into the coordinate space of the MEG data. The skull was stripped using Brain Extraction Tool, and a 3D convex hull approximating the inner surface of the skull was constructed using the software package Brainhull (http://kurage.nimh.nih.gov/meglab/Meg/Brainhull). Taking into account the position of the head relative to the sensors, a multi-sphere model (Huang et al., 1999) was computed. To normalize MEG source estimates into MNI space, we computed a nonlinear warp of each subject's brain to a single-subject template, the “colin27” brain, using the software package ANTS (Avants et al., 2011). This warp was then used to transform single-subject MEG activity maps into MNI space, and also in the reverse direction to transform virtual channel coordinate locations from MNI atlas space into individual space.

**MEG source localization**

MEG data was analyzed exclusively in source space, using synthetic aperture magnetometry (SAM, Vrba and Robinson, 2001). SAM is a scalar beamformer technique, in which activity at a single point inside the brain is reconstructed as a linear combination of all of the sensors. The weights for the linear combination are designed to pass activity originating from the point of interest while attenuating activity from everywhere else, whether of neural origin or not. Thus, analysis of source-space signals obtained by beamforming accomplishes both source localization and artifact reduction, and compensates for differences in head shape and head position across participants, facilitating multi-subject analysis. In contrast to some other beamforming techniques that reconstruct neuronal currents in 2 or 3 orthogonal directions (Huang et al., 2004), SAM chooses a single source orientation at each spatial point to maximize the variance of the obtained signal, resulting in improved signal-to-noise ratio under certain conditions (Sekihara et al., 2004).

Due to the “dual-state SAM” analysis approach using the pseudo-T algorithm to compute power differences between two conditions, multi-subject statistical maps were derived from subtractive contrast images computed on the single-subject level, not from individual conditions. Maps of pseudo-T values throughout the brain were spatially normalized to MNI space by applying the nonlinear transforms computed by ANTS (by warping the T1-weighted MRI to an MNI template), enabling random-effects analysis at the group level. SAM maps were interpolated to a voxel size of 5 mm^3^ upon warping to MNI space. Group statistics on SAM results were computed in a similar fashion as is customary in fMRI studies. For each experimental comparison, the spatially normalized whole-brain map of pseudo t-values was submitted to a voxel-wise one-sample t-test across subjects. See main text for details of the statistical inference, including correction for multiple comparisons.

**Time-frequency decomposition.**

Thirty wavelets with frequencies from 2 to 60 Hz in logarithmically spaced steps and with lengths from 3 to 10 wavelet cycles in logarithmic steps were used for time-frequency analysis.

**Stationary spectral analysis**

For each virtual channel and each trial, data from the 500-ms period of interest was subjected to multitaper spectral analysis using the MATLAB function *pmtm* with 3 Slepian tapers per trial. Spectra were averaged across trials at each channel and then averaged across participants. Because neural power spectra are dominated by a strong “red noise” or “1/f” background, we removed this background to make “whitened” spectra, on which the differences between conditions are easier to plot. The power spectra of all three conditions were averaged, and the average was fit to a function of the form

*P(f) = Af^α^*

where *f* is frequency, *P* is power, and *A* and *α* are constants fit by a nonlinear optimization program (fminsearch in matlab). The 1/f background fit was subtracted from the power spectrum of each individual condition. This whitening procedure was solely for purposes of display and did not affect any procedures of statistical inference used in this study.

**Event-related currents analysis**

Beamformer weights were computed from broadband single trial data (0-100 Hz) over the entire trial epoch time. Next, sensor signals were averaged across trials within each condition. The sensor averages were then multiplied by the beamformer weights to yield the ERCs in source space. This procedure yields a much more accurate and robust estimate of source activity compared to computing beamforming weights directly from the averaged data. One issue with the SAM procedure is that the polarity of the signal is arbitrary - the optimal current direction estimation can yield a positive current in one direction or a negative current in the opposite direction. Polarity can flip randomly between voxels and subjects. Therefore, estimated voxel-wise ERCs were rectified by converting all signal values to absolute value, and then temporally smoothed with a 9-point hamming filter. Next, we applied a normalization procedure to convert ERCs to a standardized amplitude, while maintaining relative differences between experimental conditions. This procedure was pioneered by Barca et al. (2011) to address the issue that beamformer estimates of time-domain signals can vary in amplitude by an order of magnitude or more between sites and participants, introducing the risk that participants with the largest amplitude signals will contribute disproportionally to the averages computed across participants. To correct for this, the overall mean and standard deviation was computed for the concatenation of all 8 condition time series, and each time series was individually converted into a series of z-scores using these values.

**Sensitivity power analysis**

The sample size in the present study was 20 participants, which is slightly below our usual standard of 25-30 participants for electrophysiological experiments on language. The experiment was constrained by funding and a limited period for a visiting researcher. Thus, our ability to detect subtle differences between conditions may have been limited compared to a larger study. This would especially affect the comparisons for which we observed null effects, whereas a larger study may have revealed an effect. Thus, we consider here the power achieved in the study and the minimum effect size detectable with it.

Due to the necessity to correct analyses for multiple comparisons across voxels, statistical significance in neuroimaging with cluster-level correction depends on both the effect size achieved at individual voxels and the number of adjacent voxels jointly crossing a given threshold. The voxel-level threshold of p<.01 used in this study requires larger clusters than a more stringent threshold (e.g. p<.001, common in fMRI studies), but given that source estimates of MEG activity tend to be smoother than fMRI data, large clusters are common (note that the smoothness is explicitly accounted for in the analysis programs used here, and smoother data requires still larger clusters).

Leaving aside the complexities of cluster-level correction, it is useful to examine the effect sizes obtained for the various contrasts in this experiment, considering those that did and did not achieve significance at the cluster level. A sensitivity power analysis (run in G-power 3.1.9.6; Faul et al., 2007) indicated that for our analyses, which were 2-tailed paired t-tests with n=.20 and a voxel-wise threshold of p<.01, the effect size required for 80% detection power at a single voxel would be Cohen's d = 0.84, conventionally considered to be a large effect. For a more stringent voxel-wise threshold of p<.001, we would require d = 1.08. If we were to raise the number of participants to 30, then p<.01 would require an effect size of only d = 0.66 (considered medium size), and p<.001 would require 0.83 (large). Note that an often quoted figure for "the smallest effect size of interest in psychological research" is d = 0.4, which requires over "50 participants for a simple comparison of two within-participants conditions," a standard very seldom met in neuroimaging (Brysbaert et al., 2019).

Table S1 shows the *maximum* effect sizes obtained in whole-brain maps (at any voxel) for all significant and non-significant contrasts reported in this experiment. To compute these, we measured the most significant voxel in the cluster (highest absolute value; either positive or negative) using the Z-value reported by 3dttest++ with the Clustsim option (which is based on conversion from the t-values via equivalent p-values). From this, we estimated Cohen's d with the conventional approximation

$$d=\frac{z}{\surd n}$$

| **contrast** | **figure** | **abs max Z** | **abs max d** |
| --- | --- | --- | --- |
| ERD: retention vs. intertrial | 2D | -4.88 | -1.09 |
| ERD: post vs. pre distractor | 2E | -5.46 | -1.22 |
| ERD: pho rel vs. unrel | 3A | -4.869 | -1.09 |
| ERD: sem rel vs unrrel | 3D | -3.377 | -0.76 |
| ERC: pho rel vs. unrel | 4A | 4.04 | 0.90 |
| **non-significant contrasts** |  |  |  |
| ERD: sem1 vs. sem2 | NA | -2.58 | -0.58 |
| ERD: pho1 vs. pho2 | NA | -2.94 | -0.66 |
| ERC: sem rel vs. unrel | NA | 3.91 | 0.88 |
| ERC: pho1 vs. pho2 | NA | -3.62 | -0.81 |
| ERC: sem1 vs. sem2 | NA | 3.29 | 0.74 |

Table S1. Absolute maximum Z-scores and approximately equivalent Cohen's d values for all contrasts reported in the paper, significant (with figure number) and non-significant.

Note that these are the *maximum* effect sizes observed in the maps. Effect sizes of voxels within significant clusters ranged from the individual maximum down to the threshold value at p<.01, which has a Cohen's d equivalent of 0.57. For the significant contrasts, the effect sizes are consistently large except for the weakest one, 8-30 Hz ERD for semantically related vs. unrelated distractors (figure 3D). The ERD contrasts examining position effects (relatedness to early vs. late words in the sentence) had maximum effect sizes not much higher than the threshold level of 0.57, and thus it seems likely to us that even a larger study would not demonstrate substantial position effects, although this is necessarily speculative. The effects on Event-related currents are somewhat larger. Semantically related vs. unrelated distractors yielded a maximum effect size of 0.88, which fell within a cluster of super-threshold voxels that was not large enough to be significant, but overlapped in location with the cluster seen for ERCs to phonologically related vs. unrelated distractors. Thus we may speculate that a larger study may show a significant effect for this. For position effects on event-related currents (pho1 vs. pho2, and sem1 vs. sem2), inspection of non-significant brain maps did not lead to a strong impression either way about whether a real effect may exist.
